# Precise amplification-free detection of highly structured RNA with an enhanced SCas12a assay

**DOI:** 10.1101/2024.10.20.619274

**Authors:** Junqi Zhang, Qingyuan Jiang, Wenwen Deng, Shuqi Jin, Xinping Wang, Ruyi He, Wenhao Yin, Jie Qiao, Yi Liu

**Affiliations:** State Key Laboratory of Biocatalysis and Enzyme Engineering, School of Life Sciences, Hubei University, Hubei 430042 (China); Pilot Base of Food Microbial Resources Utilization of Hubei Province, School of Life Science and Technology, Wuhan Polytechnic University, Hubei 430023 (China); State Key Laboratory of Esophageal Cancer Prevention & Treatment, Department of Oral and Maxillofacial Surgery, The First Affiliated Hospital of Zhengzhou University, Zhengzhou University, Zhengzhou 450001, China; BravoVax Co., Ltd., Wuhan, Hubei 430075 (China)

## Abstract

The CRISPR/Cas12a system has revolutionized molecular diagnostics; however, its application in directly detecting complex structured RNA remains challenging. Recently, we have developed a RNA detection method called SCas12a, which exhibits high sensitivity and efficiency in detecting RNA molecules devoid of intricate secondary structures. Here, we present an enhanced SCas12a assay (SCas12aV2) that facilitates precise amplification-free detection of highly structured RNA molecules. Our approach reengineers the split Cas12a system by optimizing the scaffold RNA length and targeting asymmetric RNA structures, thereby minimizing steric hindrance. We observe that utilization of a dsDNA-ssDNA hybrid DNA activator significantly enhances both the sensitivity and kinetics compared to those achieved using traditional ssDNA or dsDNA activators. The SCas12aV2 assay demonstrates exceptional sensitivity, with a limit of detection reaching 246 aM for pooled activators and 10 pM for single-site targeting. It also exhibits high specificity for single nucleotide polymorphisms (SNPs) and successfully identifies viable bacterial populations and SARS-CoV-2 infections from clinical samples. The assay’s versatility is further highlighted by its applicability to various Cas12a orthologs, including the thermostable CtCas12a. This work offers a significant advance in molecular diagnostics, enhancing the potential for accurate and efficient RNA detection.

## Introduction

The emergence of CRISPR (Clustered Regularly Interspaced Short Palindromic Repeats)/Cas systems has dramatically transformed molecular biology, providing exceptional precision in genome editing and regulation (1, 2). Among the diverse Cas proteins, Cas12a (3, 4) stands out as a type II-C RNA-guided DNA endonuclease. This enzyme can perform site-specific cleavage, known as *cis*-cleavage (5), on target double-stranded DNA (dsDNA). Additionally, it exhibits *trans*-cleavage activity on any single-stranded DNA (ssDNA) when a complementary sequence is identified within the spacer region of the crRNA. This unique capability has facilitated the development of highly sensitive and specific methods for nucleic acid detection, such as DETECTR (3, 6) and HOLMES (4), highlighting their potential in enhancing point-of-care diagnostic tools.

To date, Cas12a-based detection tools have predominantly targeted DNA, with the notable exception of methods incorporating reverse transcription for RNA analysis (7-9). Recently, several research groups have employed ‘split-activator’ strategies to engineer the Cas12a system for direct detection of RNA. (10-13). For example, Jain et al. developed the SAHARA diagnostic method (10), enabling programmable RNA detection with Cas12a. This technique detects RNA at picomolar concentrations by utilizing a PAM-containing dsDNA as a seed region for the crRNA. Similarly, Moon et al. developed an asymmetric CRISPR assay (12) that leverages fragmented RNA/DNA targets for miRNA detection, obviating the need for pre-amplification. Our group has recently developed a diagnostic technique known as SCas12a assay (13), which combines Cas12a with a split crRNA comprising a 20-nt scaffold RNA and a variable 20-nt spacer RNA. This method harnesses the target RNA as the spacer, achieving sensitive, amplification-free, and multiplexed RNA biomarker detection. However, all these Cas12a-based RNA detection methods encounter challenges in accurately quantifying long RNA molecules with complex secondary structures. Therefore, it is imperative to develop more robust diagnostic tools in order to overcome these limitations.

In this study, we hypothesize that reducing steric hindrance between the highly structured RNA and the DNA activator will enhance the performance of the SCas12a assay. To test this hypothesis, we modified the split Cas12a system by repositioning the target RNA further from the protospacer adjacent motif (PAM) sequence. Our findings reveal that Cas12a can accommodate various lengths of split scaffold RNAs, with a 26-nucleotide scaffold RNA (S6) being optimal for RNA detection. Additionally, we found that targeting asymmetric RNA structures and utilizing a PAM-containing dsDNA-ssDNA hybrid activator significantly improves detection efficiency and sensitivity. We designate this improved assay as SCas12aV2, which achieves a limit of detection (LoD) of 10 pM for single-site targeting and 246 aM for multi-site targeting. Furthermore, SCas12aV2 enables the effective detection of complex structures within the SARS-CoV-2 RNA target, which are inaccessible to SHERLOCK using Cas13a (14, 15). We also demonstrate the successful application of SCas12aV2 for identifying viable bacterial populations (16) and detecting SARS-CoV-2 infections in patients (17). Collectively, we have developed a highly efficient, amplification-free Cas12a-based method for the rapid, cost-effective, and accurate detection of RNA substrates with diverse sequences and complex structures.

## Results

### Cas12a tolerates different lengths of split scaffold RNA for *trans*-cleavage activity

AsCas12a is an ortholog of Cas12a that originates from *Acidaminococcus sp. BVL36*. The mature crRNA for AsCas12a is 40 nucleotides in length, consisting of a 20-nt scaffold and a complementary 20-nt spacer. Leveraging a split crRNA as illustrated in Fig. 1a, we have recently established the SCas12a assay for the direct detection of RNA molecules (13). Nonetheless, this method still faces limitations in analyzing long RNA molecules with intricate secondary structures. We hypothesized that alleviating the steric hindrance between the highly structured RNA and the DNA activator could potentially create sufficient space for accessibility of the RNA substrate, thereby overcoming these limitations. In this study, we initially optimized the design of the split Cas12a system by relocating the target RNA downstream of the PAM sequence (Fig. 1b), which would provide additional integration space. This modified system was designated as SCas12aV2.

**Fig 1.**
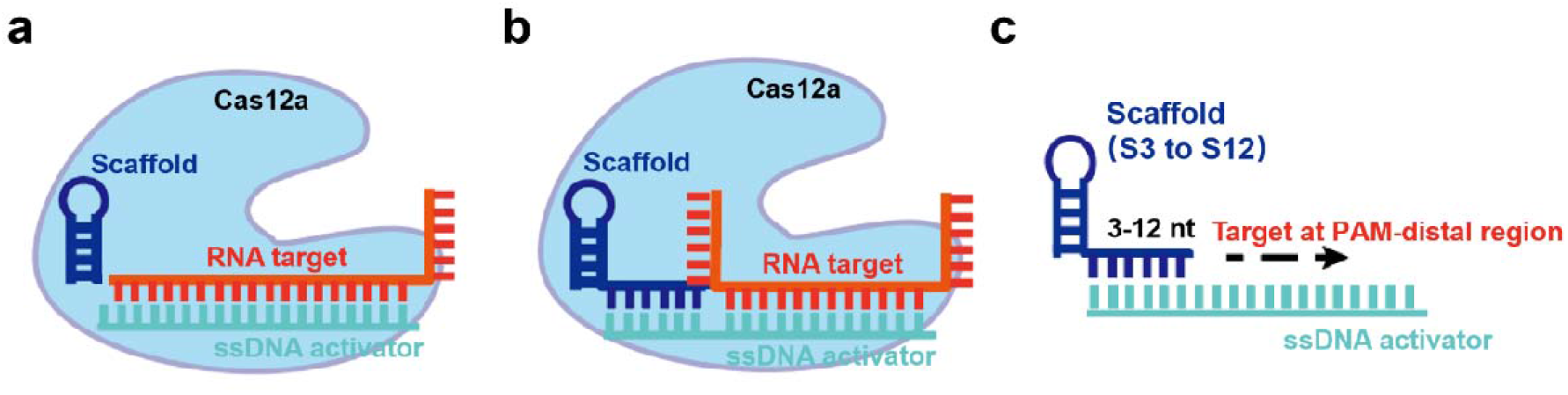
Combined use of split scaffold RNA and ssDNA activators enabling RNA detection by Cas12a. **a** Schematic illustration of the SCas12a assay by using a 20-nt scaffold RNA and ssDNA activator for RNA detection. **b** Schematic representation of Cas12a activated by combinations of scaffold RNA, ssDNA activator, and RNA target. **c** Extension of the scaffold RNA, ranging from 23 to 32 nt (S3 to S12), to detect RNA at PAM-distal region.

To activate the *trans*-cleavage activity of Cas12a, we initially assessed the viability of employing scaffold and spacer RNAs with varied nucleotide sequences. We then determined the optimal scaffold length for Cas12a by synthesizing a series of scaffold RNAs (S3 to S12), each varying in length from 23 to 32 nucleotides (Fig. 1c). For instance, using the S6 scaffold, we utilized a 14-nt spacer in conjunction with complementary target activators, which were presented as either dsDNA or ssDNA, to evaluate the enzyme’s *trans*-cleavage activity on non-specific ssDNA targets. The electrophoresis data included either a 59-bp dsDNA target (Fig. 2a) or a 59-nt ssDNA target (Fig. 2b), along with a 42-nt ssDNA reporter. Collectively, these findings demonstrate the restoration of the full *trans*-cleavage capability of the wild-type Cas12a RNP. Subsquently, the real-time fluorescence kinetic data (Fig. 2c-d) are in accordance with these PAGE results, indicating that the ssDNA activator is more effective at inducing the *trans*-cleavage activity of Cas12a. Additionally, our analysis revealed no sequence dependence for scaffold RNAs with different G-C % (Supplementary Figure S1). Based on these results, we have chosen the S6 scaffold with a consistent sequence (Table S1), for all subsequent experiments unless specified otherwise.

**Fig 2.**
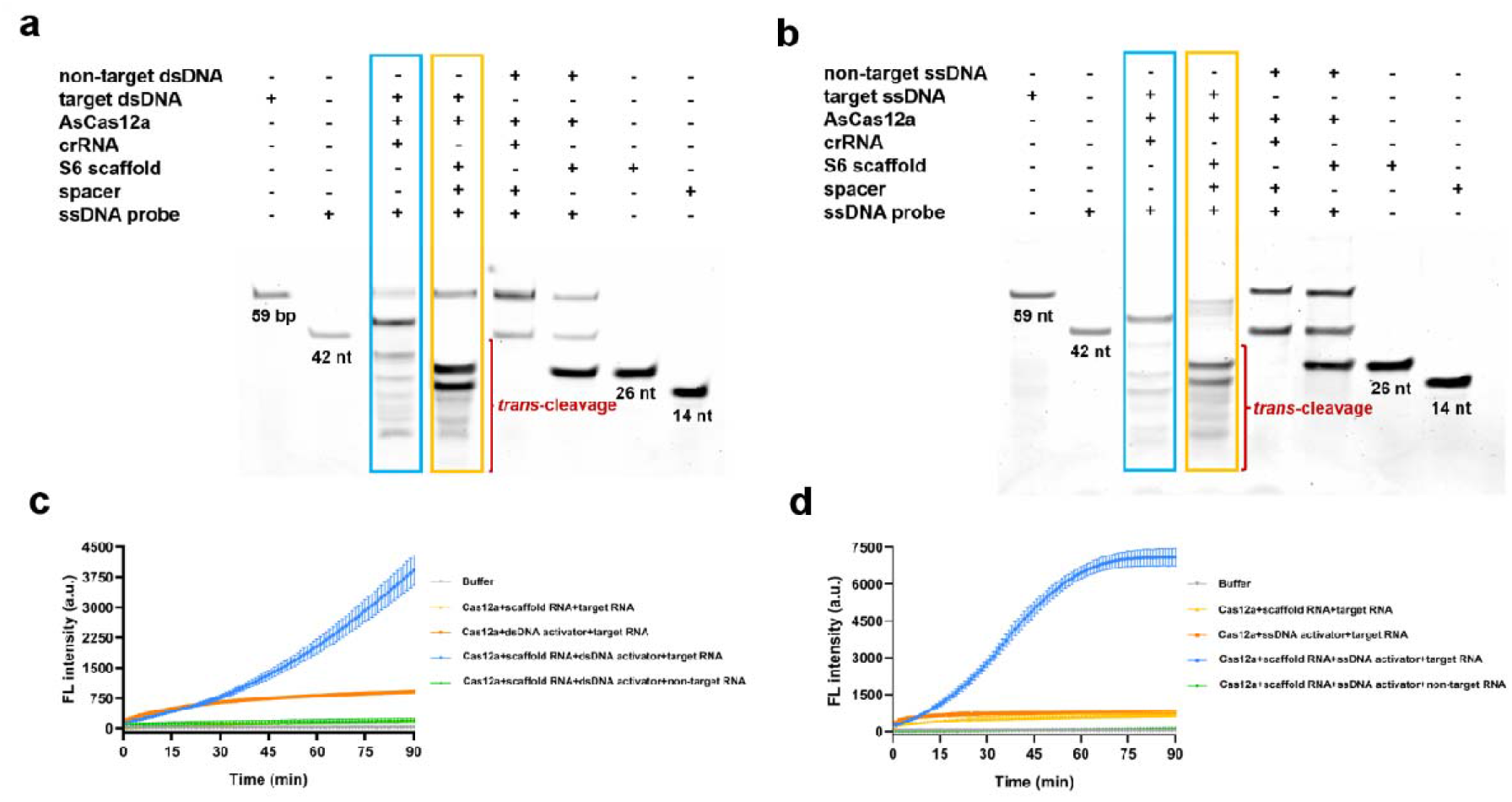
Comparison of the *trans*-cleavage activity between Cas12a with WT crRNA and split crRNA (26-nt scaffold+14-nt spacer). **a** *Trans*-cleavage of ssDNA probes by Cas12a complexed with WT crRNA (blue) and split crRNA (orange) in the presence of target or non-target dsDNA. The reactions contained 50 nM Cas12a, 100 nM of either crRNA or split RNA,10 nM of either complementary dsDNA activators or non-target dsDNA substrates, as well as 20 nM ssDNA probes. **b** *Trans*-cleavage of ssDNA probes by complexed with WT crRNA (blue) and split crRNA (orange) in the presence of target or non-target ssDNA. The reactions contained 50 nM Cas12a, 100 nM of either crRNA or split RNA, 10 nM of either complementary ssDNA activators or non-target ssDNA substrates, as well as 20 nM ssDNA probes. **c-d** Detection of target RNA by Cas12a complexed with scaffold RNA and dsDNA (c) or ssDNA (d) activators. The reaction mixtures were incubated for 60 minutes at 37°C and contained 250 nM of Cas12a, 500 nM of scaffold RNA, 10 nM target or non-target RNA substrates, 500 nM of dsDNA or ssDNA activators (dsDNA for **c** and ssDNA for **d**), and 1000 nM of fluorescent probes. For all panels, experiments were conducted in triplicate, and error bars represent the mean ± SD (n=3). Source data are provided as a Source Data file.

Next, we conducted in vitro *trans*-cleavage fluorescence assays by systematically combining a scaffold RNA with an ssDNA activator (Fig. 1c). We utilized a segment of the hepatitis C virus (HCV) polypeptide precursor RNA (Fig. 3a), characterized by its complex secondary structures, as the substrate. Specifically, we targeted four distinct regions within the HCV fragment for detection. Our findings indicate that scaffold RNAs S3 to S8 were capable of recognizing the highly structured RNA substrates (Fig. 3b-c), whereas scaffold RNAs S9 to S12 were not. Among the four target sites, S6 scaffold RNA displayed the highest detection sensitivity. Consequently, the S6 scaffold RNA was chosen for the SCas12aV2 assay. For reference, the 20-nt scaffold RNA previously employed in the SCas12a assay is denoted as S0.

**Fig 3.**
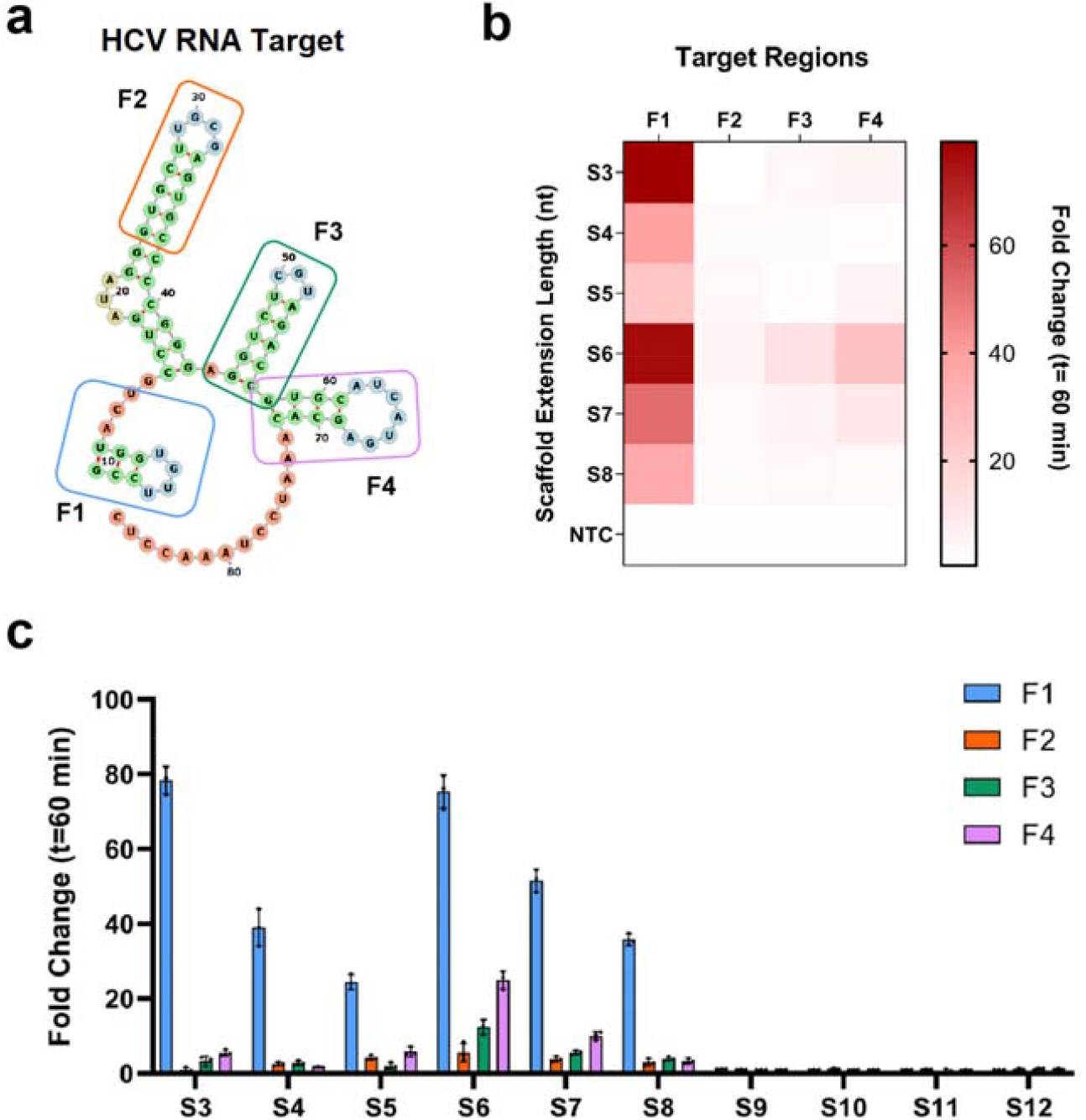
Detection of distinct regions within an HCV fragment by Cas12a complexed with S6 scaffold and ssDNA activator. **a** Schematic of an HCV RNA fragment with four target regions. The predicted secondary structure was determined by NUPACK software. **b** Heat map representing fold change with respect to NTC at t=60 min of an in vitro *trans*-cleavage fluorescence assay using the scaffold S3 to S8. **c** Comparison of the different scaffold RNAs (S3 to S12) and ssDNA activators for HCV target. The plot represents the fold change in fluorescence intensity normalized to the NTC at t=60 min (n =3).

### Enhanced sensitivity of RNA detection through recognition of asymmetric structures

The newly developed SAHARA (10) and SCas12a assay (13) enable the direct detection of RNA, yet they are limited in their ability to identify RNA substrates with intricate secondary structures, such as stems and cloverleafs. Although these methods are adept at detecting the 5’-head nucleotides, such as the F1 region in the HCV fragment, they struggle with the F2 to F4 regions which are defined by their stem structures (Fig. 3a). We hypothesized that the SCas12aV2 system could improve the accessibility of the Cas12a-scaffold RNA complex, thereby facilitating additional spatial accommodation for RNA binding. To test this, we first performed experiments targeting the symmetrical stem structures (Fig. 4a). The results demonstrated significant reductions in detection efficiency for these regions compared to F1 (Fig. 4b), likely attributable to the diminished binding affinity of the symmetrical structure towards its complementary S6 ssDNA activator. To address this issue, we then designed a series of S6 ssDNA activators specific to the asymmetric structures (Fig. 4a). The modifications resulted in significant enhancements in the fluorescence intensity (Fig. 4b), which are comparable to the levels observed for F1. Consequently, the integration of SCas12aV2 assay with asymmetric RNA targets facilitates the highly sensitive detection of intricate secondary structures within long RNA molecules.

**Fig 4.**
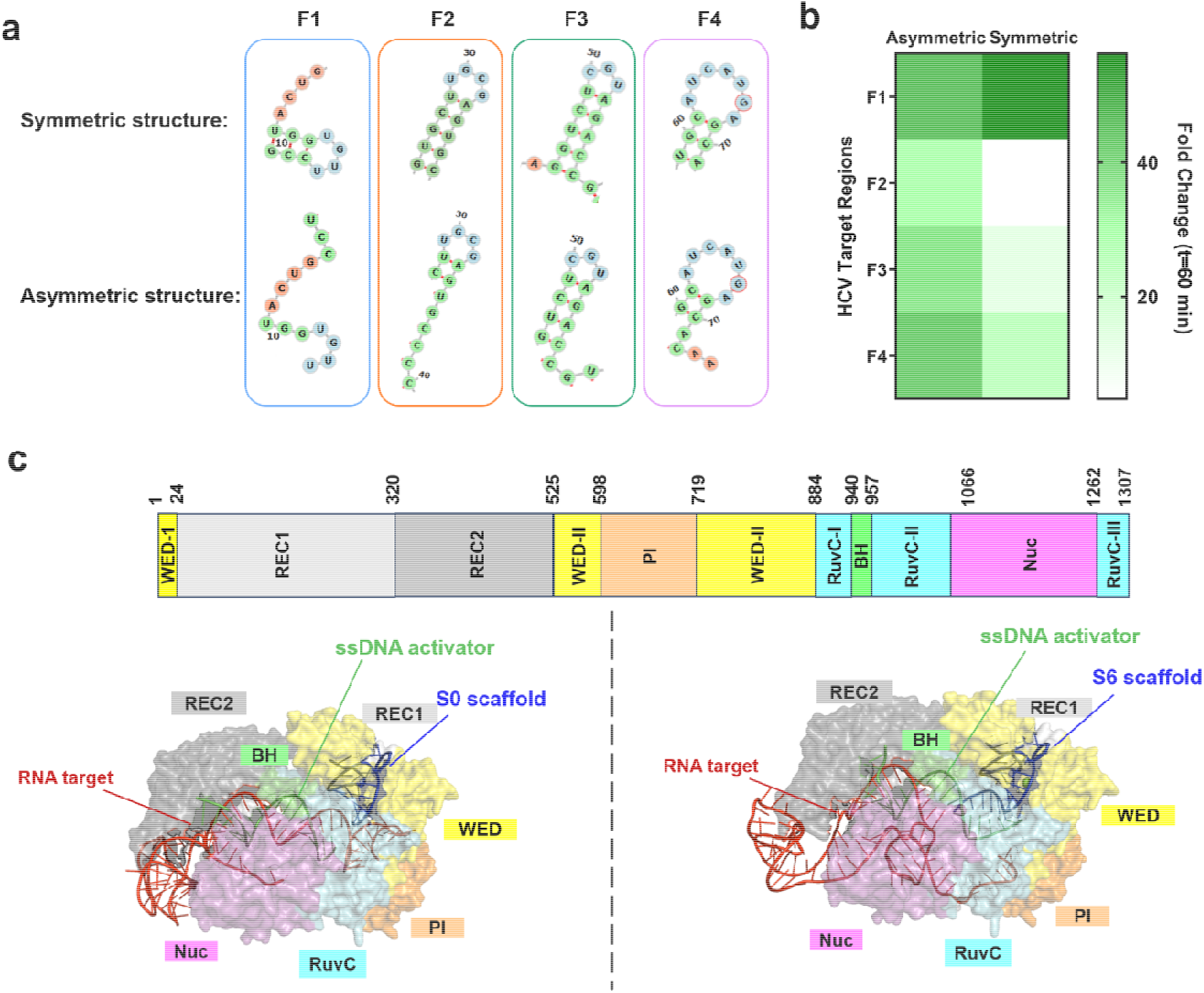
Detection of the symmetric or asymmetric structures of an HCV fragment by Cas12a complexed with S6 scaffold and ssDNA activator. **a** Schematic of an HCV RNA fragment with four target regions arranged symmetrically or asymmetrically. The predicted secondary structures were determined by NUPACK software. **b** Heat map representing fold change with respect to NTC at t=60 min of an in vitro *trans*-cleavage fluorescence assay targeting the symmetric or asymmetric structures. **c** Overall structures of the AsCas12a/S0 scaffold+ssDNA/RNA target complex and AsCas12a/S6 scaffold+ssDNA/RNA target complex. Domains are colored according to the AsCas12a (PDB: 6GQZ).

Next, leveraging the AlphaFold Server, we employed predictive structural modeling to analyze the complex comprising Cas12a, either the S0 or S6 scaffold, the ssDNA activator, and the F2 of HCV segment. The results elucidate the underlying structural basis for the heightened sensitivity observed in the SCas12aV2 assay. As illustrated in Fig. 4c, the binding of Cas12a/S6 scaffold complex to both ssDNA and RNA targets induces an activated state, thereby facilitating a highly adaptable conformation of the RNA target. The active center of Cas12a in this case exhibits flexibility, which enables efficient access of ssDNA probes to the RuvC domain for *trans*-cleavage. In contrast, the HCV target within the Cas12a/S0 scaffold complex adopts a considerably more rigid conformation, impeding effective accessibility of ssDNA probes to the RuvC domain. Recently, Huang et al (18) demonstrate that upon binding of split DNA activators adjacent to the PAM to the Cas12a RNP, a ternary complex form that can capture and interact with distal split DNA activators to achieve synergistic effects. The conclusion aligns with our experiment and predictive results, perfectly explaining the inability to detect the F2 region of the HCV target using the S0 scaffold, while detection remains feasible with the S6 scaffold.

### Enhanced capability of the SCas12aV2 assay by utilizing a dsDNA-ssDNA hybrid activator

To date, almost all detection methods based on Cas12a employ dsDNA or ssDNA as activators (Fig. 5a). We hypothesized that employing a dsDNA-ssDNA hybrid activator (Fig. 5a), capable of interacting with both the scaffold and target RNAs, might enhance the efficiency of Cas12a’s collateral cleavage activity. To distinguish this approach from the ssDNA-based method that uses S6 activator, we termed this hybrid activator as S6.1. Initially, we conducted *in vitro* collateral cleavage assays employing a scaffold RNA, a hybrid activator, and a 150-nt RNA segment originating from the human TP53 gene (Fig. 5b-h). The data revealed that the S6.1 activators displayed significantly higher fluorescence intensity compared to the S6 activators at all five targeting sites (Fig. 5g). Notably, the F4 and F5 sites, which were previously undetectable with ssDNA activators, became detectable with these hybrid activators. Furthermore, the hard-to-detect symmetrical structures at F3 and F6 were also effectively recognized, indicating that the hybrid DNA activator is compatible with a wider range of RNA substrate spatial configurations. In contrast, the SCas12a assay utilizing the S0 activator failed to detect these targets.

**Fig 5.**
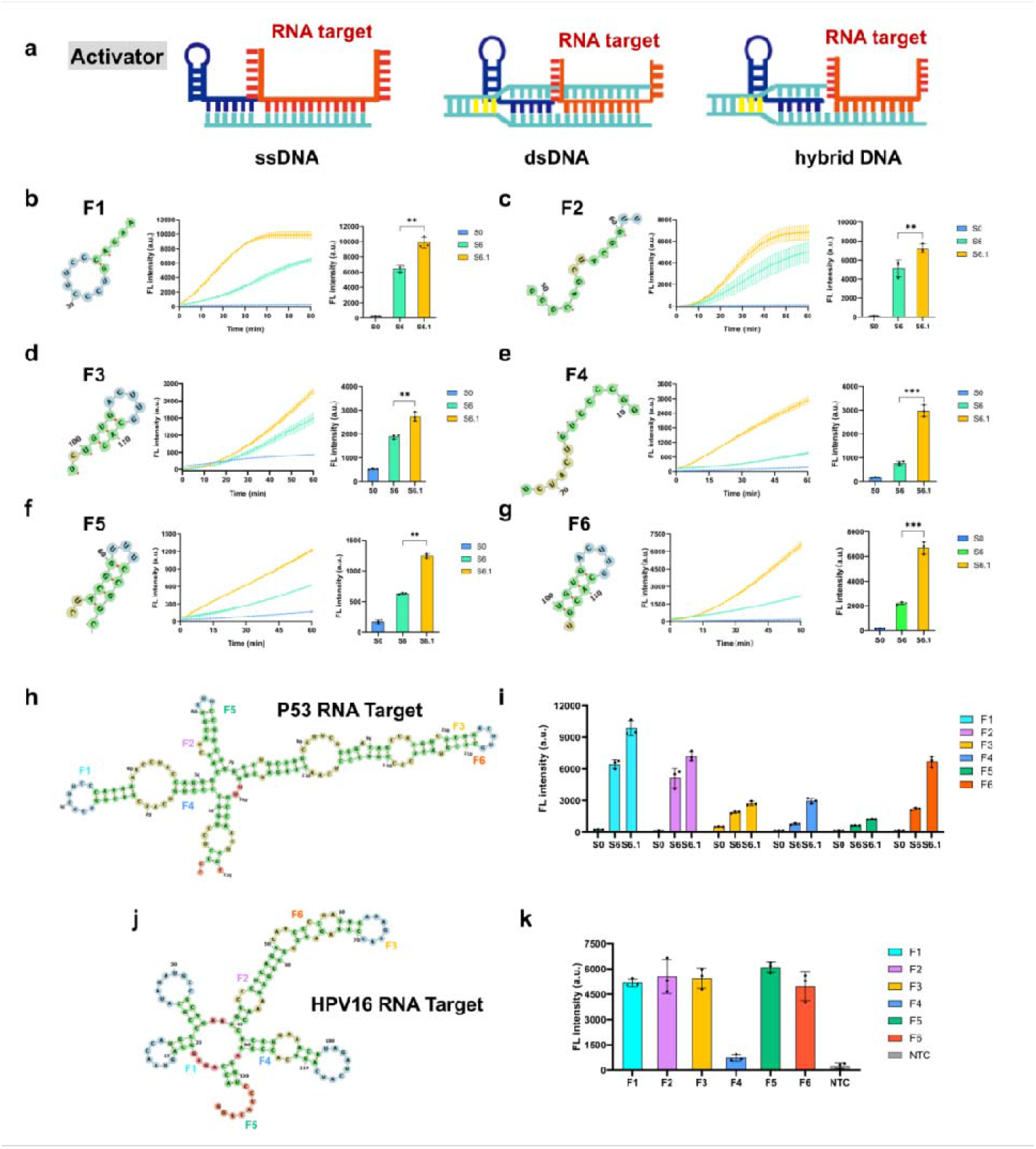
Detection of long RNA with complex structures utilizing dsDNA-ssDNA hybrid activators. **a** Schematic illustration of the SCas12a assay employing ssDNA, dsDNA, or dsDNA-ssDNA hybrid activators for RNA detection. **b-g** Comparison of the S0 (ssDNA), S6 (ssDNA) and S6.1 (dsDNA-ssDNA hybrid) DNA activators for in vitro *trans*-cleavage fluorescence assay. **h** The secondary structure of the TP53 RNA target was predicted using NUPACK software. **i** Comparative analysis of S0, S6, and S6.1 DNA activators targeting distinct regions of the TP53 RNA target. **j** The secondary structure of the TP53 RNA target was predicted using NUPACK software. **k** Comparative analysis of S0, S6, and S6.1 DNA activators targeting distinct regions of the HPV16 RNA target. For b-g, i, and k, the reaction mixtures were incubated for 60 minutes at 37°C and contained 250 nM of Cas12a enzymes, 500 nM S6 scaffold RNAs, 500 nM of DNA activators, 10 nM TP53 RNA targeted fragments and 1000 nM of fluorescent probes. All the experiments were conducted in triplicate and error bars represent mean value +/−− SD (n =3).

Next, we evaluated the kinetic properties of fluorescence assays utilizing S0, S6, or S6.1 activators. The results indicate that the catalytic reactions mediated by S6.1 are much faster than those mediated by S6 activators (Fig. 5b-g). The potential to improve the speed of point-of-care testing (POCT) for RNA molecules, especially those with long lengths and complex secondary structures, is highlighted by this observation. To confirm the method’s universality, we used the SCas12aV2 assay with S6.1 activators to detect various RNA substrates. The transcript from the human papillomavirus (HPV), known for its complex secondary structures (Fig. 5j), was tested and showed detectability of all targets except F4 (Fig. 5k). Finally, we demonstrate that SCas12aV2 facilitates the efficient detection of complex regions within the SARS-CoV-2 target (Supplementary Figure S2), which are inaccessible to Cas13a-based SHERLOCK method (17).

In conclusion, the dsDNA-ssDNA hybrid activator exhibits a significant enhancement in the performance of the fluorescence assay compared to traditional ssDNA activators. Therefore, unless otherwise specified in subsequent sections, this type of hybrid activator is adopted. Furthermore, the results demonstrate the potential of SCas12aV2 assay as a robust RNA detection tool.

### Application of the SCas12aV2 assay to different Cas12a orthologs

The orthologs of Cas12a, LbCas12a (19) and CtCas12a (20), derived from *Lachnospiraceae bacterium ND2006* and *Clostridium thermobutyricum* respectively, are referred to as Lb and Ct in this context. To date, AsCas12a and LbCas12a have seen widespread application in nucleic acid detection. Of particular interest is the recently discovered thermostable CtCas12a, which exhibits activity over a broad temperature range (20), extending from 17 to 77^°^C. Our investigation focused on assessing the performance of these Cas12a orthologs in the SCas12aV2 assay. The results demonstrate that the hybrid dsDNA-ssDNA activator is optimal for engaging all Cas12a orthologs with TP53 RNA targets (Fig. 6). Specifically, LbCas12a exhibits a significantly reduced detection efficiency, approximately 10-fold lower than AsCas12a and 20-fold lower than CtCas12a. This suggests a stringent specificity of LbCas12a’s catalytic active site towards the structural features of RNA substrates. Additionally, we observed that CtCas12a displays enhanced sensitivity in detecting RNA molecules with intricate structures and high GC content when operated at its optimal reaction temperature of 55^°^C. These observations suggest that the selection of Cas12a orthologs can be customized to meet specific requirements within the SCas12aV2 assay framework in the future.

**Fig 6.**
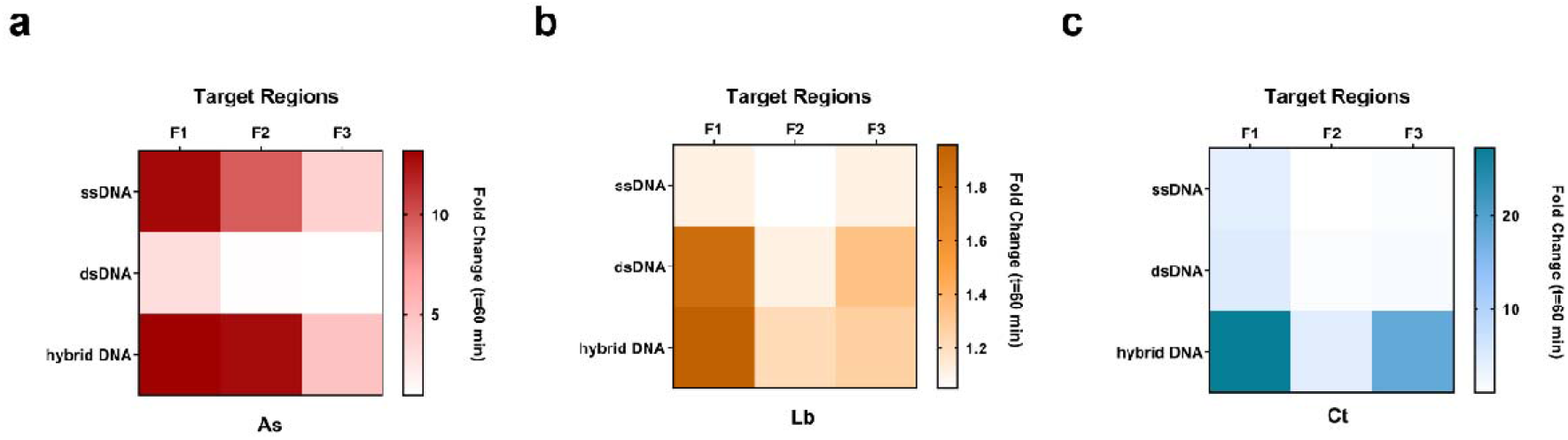
Validating the universality of SCas12aV2 assay by utilizing Cas12a orthologs. **a-c** Heat maps representing the fold changes in the fluorescence intensity of in vitro trans-cleavage assay (n=3) with Cas12a orthologs. The reactions contained 250 nM of As, Lb, or Ct Cas12a enzymes, 500 nM S6 scaffold RNAs, 500 nM of ssDNA, dsDNA, or dsDNA-ssDNA hybrid activators, 10 nM TP53 RNA targeted fragments and 1000 nM of fluorescent probes. All the experiments were conducted in triplicate and error bars represent mean value +/−− SD (n =3).

### The limit of detection for RNA detection by SCas12aV2 assay

We first employed the fluorescence-based SCas12aV2 assay for RNA detection without the need for amplification. The sensitivity was evaluated by employing a panel of dsDNA-ssDNA hybrid activators targeting specific RNA segments derived from human immunodeficiency virus (HIV) (Fig. 7a) and TP53 (Supplementary Figure S3). Regarding the HIV RNA target, we generated a gradient of RNA target concentrations ranging from 246 aM to 10 nM. The limit of detection (LoD) for each target was determined by introducing an excess amount of these activators, resulting in an established LoD of 100 fM as described in detail in the Methods section. For the TP53 RNA targets, distinct regions exhibited varying LoDs ranging from 3.6 pM to 86 pM (Supplementary Figure S3). Overall, the average LoD of 10 pM demonstrates the assay’s efficacy in detecting RNA with diverse secondary structures. Furthermore, we have successfully established a linear correlation between the fluorescence intensity and the concentration of TP53 RNA standards (Supplementary Figure S4). This validation confirms the robustness of our methodology for quantifying RNA. Moreover, employing a combination of activators that target multiple regions of the identified RNA enables a significant enhancement in LoD by 100-to 1000-fold. For instance, we employed a panel of activators targeting four distinct regions within the RNA of SARS-CoV-2 virus (Supplementary Figure S5), resulting in an estimated LoD value of 246 aM. Finally, we demonstrate that SCas12aV2 facilitates the efficient detection of complex regions within the SARS-CoV-2 target, which are inaccessible to Cas13a-based SHERLOCK method (17).

**Fig 7.**
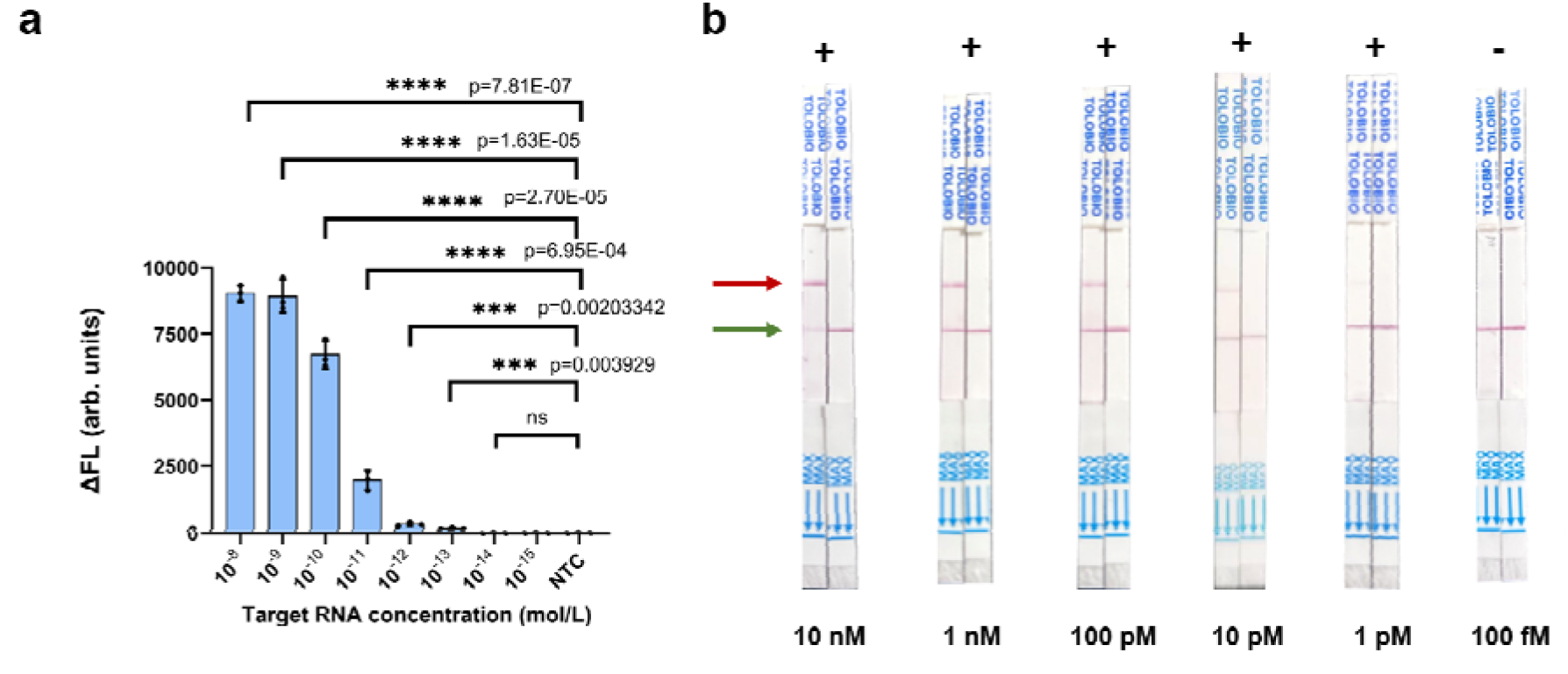
Determination of the limit of detection of the SCas12aV2 assay for RNA detection. **a** Limit of detection of the HIV target was determined by a fluorescence assay. The plot illustrates the background-subtracted fluorescence intensity at t = 60 minutes for varying concentrations of the target. All the experiments were conducted in triplicate and error bars represent mean value +/−− SD (n =3), and statistical analysis was conducted using a two-tailed t-test. Statistical significance was determined as follows: ns (not significant) for p > 0.05, ^*^ for p ≤ 0.05, ^**^ for p ≤ 0.01, ^***^ for p ≤ 0.001, and ^****^ for p ≤ 0.0001. **b** Limit of detection of the HIV target was determined by a lateral flow assay. Similar experiments were conducted, with the exception that a dipstick reporter was employed following a 60-minute LFA cleavage reaction. The red arrow indicates the test bands, while the green arrow indicates the control bands. The results are denoted by the symbols “+” for positive outcomes and “-” for negative outcomes. Source data are provided as a Source Data file.

In addition to fluorescence readout, we also assessed the sensitivity of SCas12aV2 using a rapid lateral flow assay (LFA) readout, achieving an impressive limit of detection (LoD) as low as 1 pM (Fig. 7b). Collectively, these experiments validate the efficient and unbiased capability of the SCas12aV2 assay in detecting RNA targets.

### SCas12aV2 assay exhibits exceptional mutation detection specificity

In our recent research, we established that the SCas12a assay possesses remarkable sensitivity for the detection of single nucleotide polymorphisms (SNPs) within miRNAs (13). Based on these findings, the current study aimed to evaluate the sensitivity of the SCas12aV2 assay for SNP detection in long RNA molecules. Utilizing an HIV-derived fragment as a case study, we engineered 14-nt ssDNA activators (S6) and dsDNA-ssDNA DNA hybrid activators (S6.1) with single-point mutations for SCas12aV2 detection. For comparative analysis, we also designed 20-nt ssDNA activators with corresponding mutations for use with the DETECTR, employing wild-type crRNA (Fig. 8). The specificity of target recognition was ascertained by quantifying the relative fluorescence enhancement between mismatched and perfectly matched targets. A diminished relative fluorescence enhancement rate indicates enhanced specificity, as it suggests a greater capacity to distinguish between mismatched and matched targets. Our findings reveal that the detection sensitivity for single-point mutations was position-dependent in both the SCas12aV2 assay and the DETECTR system when compared to the wild-type target. Notably, the S6 ssDNA activator outperformed the S6.1 dsDNA-ssDNA hybrid in SNP detection. Specifically, mutations at positions M4, M6-9, and M10 substantially reduced the SCas12aV2-mediated *trans*-cleavage activity. In addition, our data highlight that SCas12aV2 exhibits enhanced specificity for mutations located in PAM-distal regions. Together, these results underscore the SCas12aV2 assay’s exceptional reliability in detecting SNPs within RNA targets.

**Fig 8.**
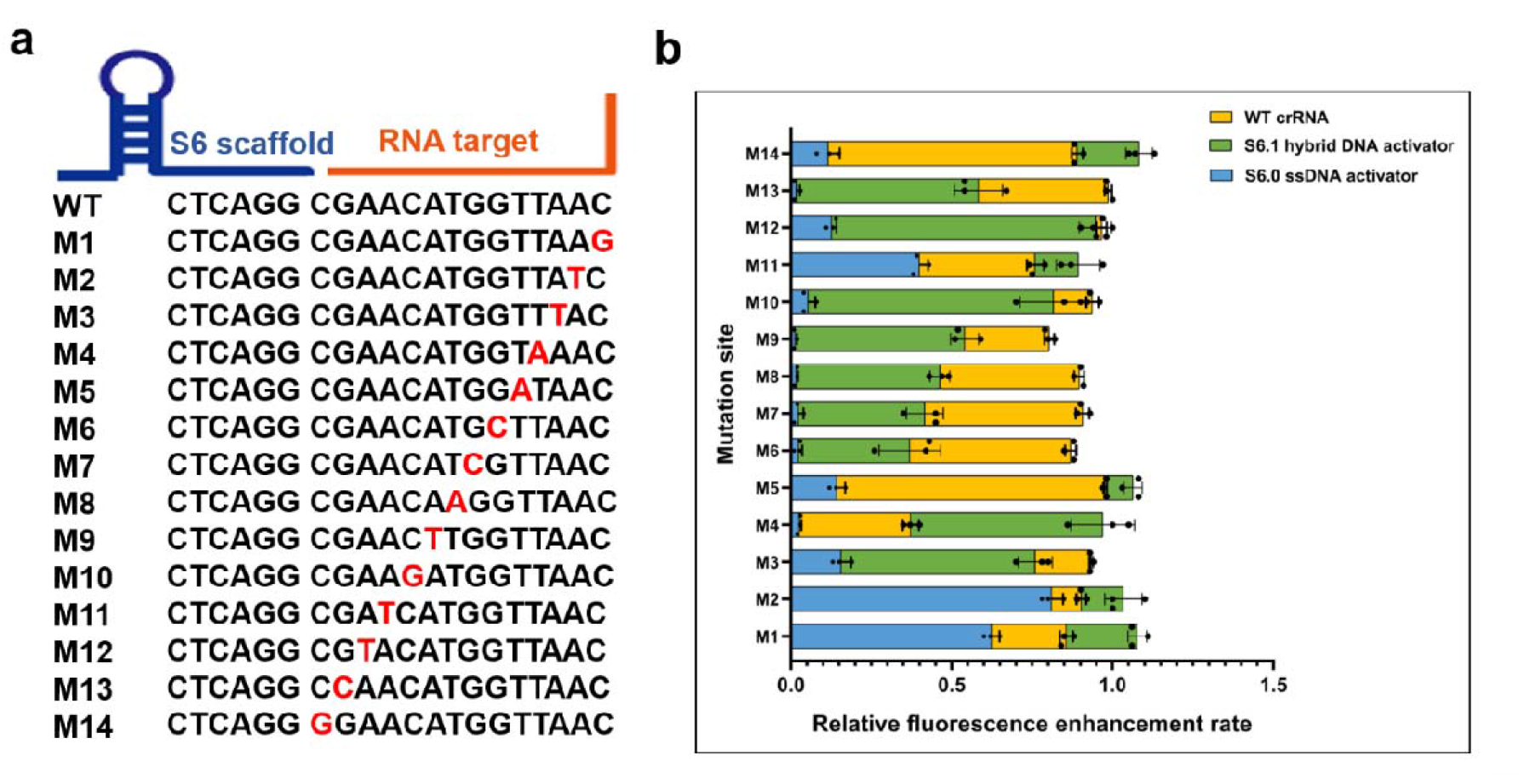
Specificity of SCas12aV2 assay towards single point mutations in target. **a** ssDNA activators were designed with point mutations across the length of the pairing region in an HIV target. The mutation location is identified by ‘M’ following the nucleotide number where the base has been changed to its complementary nucleotide (3’ to 5’ direction). **b** Comparison of fluorescence fold changes in the *trans*-cleavage assay between DETECTR and SCas12aV2 utilizing either S6 ssDNA or S6.1 hybrid DNA activators. All fluorescence values were normalized to those of WT activator. Statistical analysis for n=3 biologically independent replicates comparing the normalized fold change for DETECTR vs SCas12aV2 assay. Statistical analysis was conducted using a two-tailed t-test. Statistical significance was determined as follows: ns (not significant) for p > 0.05, ^*^ for p ≤ 0.05, ^**^ for p ≤ 0.01, ^***^ for p ≤ 0.001, and ^****^ for p ≤ 0.0001. error bars represent mean value +/−− SD (n =3). Source data are provided as a Source Data file.

### Application of the SCas12aV2 assay for practical studies

The accurate clinical identification of pathogen-induced infections necessitates the utilization of highly sensitive and rapid RNA detection methods. In this study, we validated the effectiveness of the SCas12aV2 assay in detecting viable bacterial populations ranging from 0% to 100%, as well as in clinical specimens from individuals infected with SARS-CoV-2. Figure 9a depicts the preparation of bacterial suspensions containing varying proportions of viable and dead ATCC25922 *E. coli* within a volume of 1 mL. The ratio between viable and total bacteria was systematically adjusted from 0% to 100%. Initially, traditional plate counting techniques were employed to assess the viability of these samples by correlating colony counts with viable bacterial cell content (Fig. 9b). Subsequently, RNA extracted from these samples underwent SCas12aV2 analysis. A direct correlation was observed between fluorescence intensity and the proportion of viable bacteria (Fig. 9c). These findings demonstrate that the SCas12aV2 assay is capable of quantifying viable bacterial counts, exhibiting an enhanced ability to detect viable bacteria as the ratio between viable and total bacteria increases. Finally, we successfully isolated five RNA samples from nasal swab specimens collected from individuals confirmed to have SARS-CoV-2. The clinical RT-qPCR analysis yielded Ct values ranging from 10.87 to 22.23 for these samples. Employing the SCas12aV2 direct detection assay, which targets four distinct regions of the SARS-CoV-2 genome, we accurately identified all five samples as positive, with a clear distinction in slopes compared to the negative control swab depicted in Fig. 9e. The slopes of the positive samples were significantly correlated with their respective input concentrations, underscoring the assay’s quantitative capability. The inclusion of a negative swab control enabled us to mitigate the impact of matrix effects inherent in clinical specimens. Additionally, we conducted tests with a non-targeting control S6.1 activator on all samples, which yielded no significant signals.

**Fig 9.**
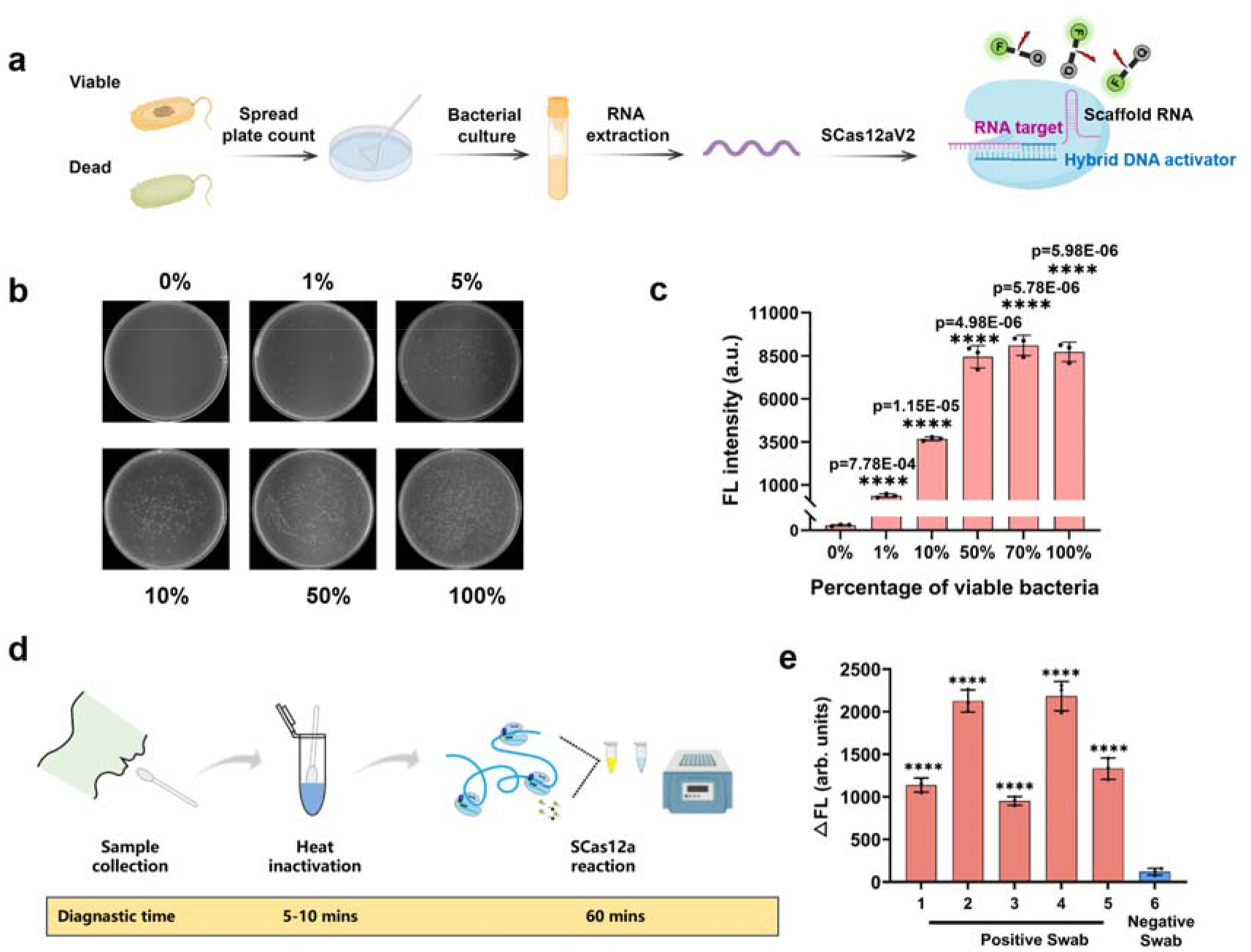
Utilization of the SCas12aV2 assay for pathogen detection. **a** Schematic diagram of differentiating ATCC25922 *E. coli* viable and dead bacteria by SCas12aV2 assay. **b** The ratio of viable bacteria to total bacteria was verified using standard plate counts as 0, 1, 5, 10, 50 and 100%, respectively. **c** Analyzing viable pathogenic bacteria using the SCas12aV2 assay. The fluorescence response of SCas12aV2 was assessed for profiling bacteria at various percentages of viable bacteria (0, 1, 5, 10, 50, and 100%). The reaction employed *E. coli* 16S rRNA primer1, 1 μM *E. coli* 16S rRNA primer2, and *E. coli* 16S rRNA targeted S6.1 activator, respectively. **d** Schematic diagram of detecting SARS-Cov-2 from the clinical swab samples by SCas12aV2 assay. **e** Pre-extracted RNA from five nasopharyngeal swabs confirmed positive for SARS-CoV-2 by RT-qPCR was tested by SCas12aV2 assay. A confirmed negative swab was tested for comparison. All fluorescence values were normalized to those of WT activator. Statistical analysis for n=3 biologically independent replicates comparing the normalized fold change for DETECTR vs SCas12aV2 assay. Statistical analysis was conducted using a two-tailed t-test. Statistical significance was determined as follows: ns (not significant) for p > 0.05, ^*^ for p ≤ 0.05, ^**^ for p ≤ 0.01, ^***^ for p ≤ 0.001, and ^****^ for p ≤ 0.0001. error bars represent mean value +/−− SD (n =3). Source data are provided as a Source Data file.

## Discussion

This study introduces an innovative enhancement to the CRISPR/Cas12a-based detection system, specifically the SCas12aV2 assay, which holds significant potential for the precise detection of highly structured RNA molecules. The key advantages of the SCas12aV2 assay are as follows: (1) it enables direct detection of highly structured RNA molecules without amplification, simplifying the process and reducing the risk of contamination and false positives; (2) it minimizes steric hindrance through structure-based design, including optimization of scaffold RNA length, targeting asymmetric RNA structures, and adopting a dsDNA-ssDNA hybrid as the activator to initiate *trans*-cleavage activity; (3) it is applicable to various Cas12 orthologs, including the thermostable CtCas12a, highlighting the assay’s versatility and suitability for diverse conditions. Moreover, this method exhibits high sensitivity, achieving a LoD as low as 246 aM when using pooled DNA activators. Its specificity for SNPs confirms the assay’s reliability in detecting SNPs within RNA targets. Additionally, the SCas12aV2 method accurately detects viable bacterial populations and RNA from clinical specimens, as evidenced by its application to SARS-CoV-2.

While the SCas12aV2 assay presents notable improvements, it faces certain challenges. A significant limitation is the reliance on the design of activators for each target RNA, a process that can be time-consuming and may necessitate substantial optimization. Additionally, the assay, although highly sensitive, may encounter difficulties in distinguishing between very similar sequences, potentially affecting its utility in diagnostic applications where sequence specificity is critical. To overcome these limitations, future refinements of the SCas12aV2 assay could integrate machine learning algorithms to predict and optimize activator designs, thereby streamlining the development process and reducing the time required for assay preparation.

In conclusion, the SCas12aV2 assay constitutes a substantial advancement in molecular diagnostics, providing a sensitive, specific, and user-friendly method for the detection of structured RNA molecules. Its broad potential applications in research, clinical diagnostics, and infectious disease monitoring are evident. The “split-activator” system design has been validated for use with not only Cas12a orthologs but also other CRISPR/Cas enzymes, such as Cas13 and Cas14. We foresee the development of increasingly potent Cas enzyme-based detection tools in the future, advancing from the discoveries reported in this manuscript.

## Materials and Methods

### Ethical statement

For this study, human nasal swab specimens for SARS-CoV-2 detection were collected and provided by College of Medicine and Health Science with a protocol approved by the ethics committee at Wuhan Polytechnic University.

### Materials

Fluorescence-based detection reagents, including DNA, RNA, FAM-tagged ssDNA, and FAM-tagged RNA, were sourced from Sangong Biotech (Shanghai, China) as detailed in Supplementary Table S1-S3. Buffer reagents were procured from Sinopharm (Beijing, China). Enzymatic reactions were facilitated by the use of 10 x NEB buffer 2.1 and 10 x Cutsmart buffer, both of which were supplied by New England Biolabs (Beijing, China). The enzymatic degradation of proteins was carried out using proteinase K, and total RNA was isolated using a kit, both products being from Takara (Dalian, China). The experiments were conducted using ultrapure water to ensure the integrity of the reactions. Prior to use, all DNA and RNA molecules were resuspended in DEPC-treated water to eliminate RNase activity and then preserved at -20°C for future experimental procedures.

### Recombinant Cas12a orthologs and LwaCas13a protein expression and purification

Synthetic genes encoding the CRISPR-associated proteins AsCas12a, LbCas12a, CtCas12a, and LwaCas13a were cloned into the pET-28a(+) expression vector to construcr plasmids for recombinant protein production. The sequence accuracy of these plasmids was verified prior to transformation into *Escherichia coli* BL21 (DE3) competent cells. For expression, a single colony was inoculated into LB broth supplemented with ampicillin (100 μg/mL) and incubated overnight. The following day, these cultures were diluted into 1-liter Terrific Broth (TB) medium to an optical density at 600 nm (OD_600_) of 0.8, cooled on ice for 10 minutes, induced with 0.5 mM isopropyl β-D-1-thiogalactopyranoside (IPTG), and further incubated at 18°C for 16 hours. The bacterial cells were then pelleted by centrifugation and lysed in buffer A (20 mM Tris-HCl, pH 7.5, 300 mM NaCl, 1 mM phenylmethanesulfonyl fluoride (PMSF), 5 mM β-mercaptoethanol) containing a cocktail of proteinase inhibitors. The recombinant Cas proteins were purified using immobilized metal affinity chromatography (IMAC) with Ni-NTA resin, followed by size-exclusion chromatography on a HiLoad® 16/600 Superdex® column as per a published protocol (21). The purified proteins were concentrated using a 100 kDa molecular weight cut-off (MWCO) centrifugal filter and their concentrations determined by the Bradford assay. Subsequently, the concentrated proteins were flash-frozen in liquid nitrogen and stored at -80°C for subsequent applications.

### Gel electrophoresis

Reconstitution of wild-type Cas12a and SCas12aV2 ribonucleoproteins was achieved by mixing the respective Cas12a nucleases with either full-length crRNA or split crRNA at a 1:2 molar ratio in a reaction buffer, followed by a 20-minute incubation at ambient temperature. To assess the *trans*-cleavage activity of both wild-type Cas12a and SCas12aV2 RNPs, experiments were conducted under identical conditions as previously detailed (21), with the exception of the inclusion of a 20 nM concentration of a 42-nt ssDNA reporter. This reporter was mixed with either 10 nM of 59-bp dsDNA targets (Fig. 2a) or 59-nt ssDNA targets (Fig. 2b). The reaction outcomes were resolved on a 12% polyacrylamide gel, utilizing 1X TBE buffer for electrophoresis at a steady voltage of 100 V over a period of 2 hours. Subsequently, the gel was visualized by staining with GoldView nucleic acid stain and examined under UV illumination for documentation.

### Application of the SCas12aV2 fluorescence assay for RNA detection

Fluorescence-based detection assays were performed using a CFX96 touch real-time PCR system (Bio-Rad, Hercules, CA, USA). The assays began with the preparation of a reaction mixture consisting of 250 nM Cas12a, 500 nM scaffold RNA, and 500 nM of either ssDNA, dsDNA, or dsDNA-ssDNA hybrid activators. The components were mixed in NEB buffer 2.1 and nuclease-free water, followed by a 10-minute incubation at room temperature to allow equilibration. Next, 1000 nM FQ reporter and the appropriate concentration of target RNA were added to each reaction, bringing the total volume to 20 µl. The reactions were then incubated at 37°C for 60 minutes, monitoring the fluorescence emission at 520 nm. For LwaCas13a-mediated assays, the protocol was identical, except for the use of a FAM-ssRNA reporter.

In parallel, lateral flow assays were conducted with a dipstick containing ssDNA probes for detection. The LFA dipsticks, obtained from ToloBio (Shanghai, China), incorporated a FAM-biotin reporter for immunochromatographic analysis. In negative samples, the gold particle-biotin antibody complex bound to the FAM-biotin reporter, which was captured by the anti-FAM antibody on the test strip. Positive samples, however, exhibited cleavage of the FAM-biotin reporter by active Cas12a, resulting in a visible concentration of the gold particle-biotin antibody complex on the test line and a diminished signal on the control line.

### RNA extraction and RT-qPCR for detection of RNA

RNA was extracted from ATCC25922 *E. coli* and SARS-CoV-2 lysis via the RNA extraction kit from GenePharma (Suzhou, China). Next, the cDNA was synthesized from the extracted total RNA using the miRCURY LNA RT Kit (Qiagen). Briefly, a reverse transcription reaction was performed at 42°C for 60 min and then inactivated at 95°C for 5 min. Synthesized cDNA was stored at -20°C before use. For detection of specific RNA targets, the primers and RT-qPCR detection kit were designed and ordered from GenePharma (Suzhou, China). Subsequently, the RNA targets in the biological samples were quantitatively assessed according to the protocol provided by the manufacturer. The real-time quantitative polymerase chain reaction (RT-qPCR) reactions were performed using a CFX96 touch real-time PCR system (Bio-Rad, CA, USA) and the RNA targets were identified utilizing the SYBR green fluorescent method. The thermal cycling conditions included an initial denaturation step at 94°C for 3 minutes, followed by 40 cycles of denaturation at 94°C for 12 seconds and annealing/extension at 62°C for 30 seconds.

### Limit of detection calculation

The limit of detection (LoD) was determined by conducting the trans-cleavage assay using various dilutions of RNA or DNA targets. The value of LOD was calculated by using formula 3σ/slope, where σ is the standard deviation of three blank solutions.

### Statistics and reproducibility

No statistical method was used to predetermine sample size. No data were excluded from the analyses. The experiments were not randomized. The Investigators were not blinded to allocation during experiments and outcome assessment.

### Reporting summary

Further information on research design is available in the Nature Portfolio Reporting Summary linked to this article.

## Data Availability

All the data supporting the findings of this study are available within the Article and Supplementary Files. Source data is available in the Source Data file. Source data are provided with this paper.

## Acknowledgments

This work was supported by the National Key Research and Development Program of China 2022YFC2304304 (Y. L.), Science and Technology Innovation Talent Plan of Hubei Province 2023DJC136 (Y. L.), the Open Funding Project of the State Key Laboratory of Esophageal Cancer Prevention K2022-008 (J. Q.), the Open Funding Project of the State Key Laboratory of Biocatalysis and Enzyme Engineering SKLBEE2022022 (J. Q.), Research Funding of Wuhan Polytechnic University NO.2022RZ031 (J. Q.).

## Author Contributions

J. Q., J. Z., and Y. L. designed research; J. Q., Z. J., X. W., J. Z., Q. J., and W. Y. performed research; R. H. collected the patient samples; Y. C., J. Z., Q. J., and Y. L. analyzed data; and Y. L. wrote the paper.

