## Supplementary material for "Precise amplification-free detection of highly structured RNA with an enhanced SCas12a assay": Supplemenral files

**This PDF file includes:**

Table S1-S3

Fig. S1 to S5

**Table S1:** List of the split crRNA (5’→3’) used in this study .

| Name | Sequence |
| --- | --- |
| S0 | AAUUUCUACUAAGUGUAGAU |
| S3 | AAUUUCUACUAAGUGUAGAU GAG |
| S4 | AAUUUCUACUAAGUGUAGAU GAGU |
| S5 | AAUUUCUACUAAGUGUAGAU GAGUC |
| S6 | AAUUUCUACUAAGUGUAGAU GAGUCC |
| S7 | AAUUUCUACUAAGUGUAGAU GAGUCCC |
| S8 | AAUUUCUACUAAGUGUAGAU GAGUCCCG |
| S9 | AAUUUCUACUAAGUGUAGAU GAGUCCCGC |
| S10 | AAUUUCUACUAAGUGUAGAU GAGUCCCGCC |
| S11 | AAUUUCUACUAAGUGUAGAU GAGUCCCGCCU |
| S12 | AAUUUCUACUAAGUGUAGAU GAGUCCCGCCUC |

**Table S2:** List of the DNA activators (5’→3’) used in this study .

| Universal primer | S6.1-F | CGACGCTCTCCCTTATTTAGAGTCC |
| --- | --- | --- |
| HIV | S6-F1 | CAATAGCAATTGGTACAAGC |
|  | S6-F2 | ATGAAAGCAACACTTTTTAC |
| HCV | S6-F1 | CAGTACCACAAGGCGGACTC |
|  | S6-F2 | GCACTCGCAAGCACGGACTC |
|  | S6-F3 | CGGTCTACGAGACCTGGACTC |
|  | S6-F4 | TGCTCATGATGCACGGACTC |
|  | S6-F1 (asymmetric) | AGGCAGTACCACAAGGACTC |
|  | S6-F2 (asymmetric) | GGGGCACTCGCAAGGGACTC |
|  | S6-F3 (asymmetric) | ACGGTCTACGAGACGGACTC |
|  | S6-F4 (asymmetric) | TTGTGCTCATGATGCGGACTC |
| P53 | S6-F1 | TTCTGGGAAGGGACGGACTC |
|  | S6-F2 | AACCGTAGCTGCCCGGACTC |
|  | S6-F3 | GTGCAAGTCACAGAGGACTC |
|  | S6-F4 | AGATGACAGGGGCCGGACTC |
|  | S6-F5 | GACGGAAACCGTAGGGACTC |
|  | S6-F6 | ACGTGCAAGTCACAGGACTC |
|  | S6.1-F1 | TTCTGGGAAGGGACGGACTCTAAATAAGGGAGAGCGTCG |
|  | S6.1-F2 | GGGCAGCTACGGTTGGACTCTAAATAAGGGAGAGCGTCG |
|  | S6.1-F3 | GTGCAAGTCACAGAGGACTCTAAATAAGGGAGAGCGTCG |
|  | S6.1-F4 | AGATGACAGGGGCCGGACTCTAAATAAGGGAGAGCGTCG |
|  | S6.1-F5 | GACGGAAACCGTAGGGACTCTAAATAAGGGAGAGCGTCG |
|  | S6.1-F6 | ACGTGCAAGTCACAGGACTCTAAATAAGGGAGAGCGTCG |
|  | S6.2-F1 | GTGGTGGAATTCTGCAGATTTAGAGTCCGTCCCTTCCCAGAA |
|  | S6.2-F2 | GTGGTGGAATTCTGCAGATTTAGAGTCCGGGCAGCTACGGTT |
|  | S6.2-F3 | GTGCAAGTCACAGAGGACTCTAAATCTGCAGAATTCCACCAC |
| miRNA21 | S6 | AGTCTGATAAGCTAGGACTC |
|  | S6.1 | AGTCTGATAAGCTAGGACTCTAAATAAGGGAGAGCGTCG |
| SARS-CoV-2 | S6.1-F1 | CGGGGTGCATTTCGGGACTCTAAATAAGGGAGAGCGTCG |
|  | S6.1-F10 | GCCATTCTAGCAGGGGACTCTAAATAAGGGAGAGCGTCG |
|  | S6.1-F12 | TTACCAGACATTTTGGACTCTAAATAAGGGAGAGCGTCG |
|  | S6.1-L-F4 | CAAGACGCAGTATTGGACTCTAAATAAGGGAGAGCGTCG |
|  | S6.1-L-F5 | TGCCATGTTGAGTGGGACTCTAAATAAGGGAGAGCGTCG |
|  | S6.1-L-F6 | GTCCTCGAGGGAATGGACTCTAAATAAGGGAGAGCGTCG |
|  | S6.1-L-F14 | CCAGACATTTTGCTGGACTCTAAATAAGGGAGAGCGTCG |
|  | S6.1-L-F16 | TAGTTCCTGGTCCCGGACTCTAAATAAGGGAGAGCGTCG |
|  | S6.1-L-F17 | AGTTCCTTGTCTGAGGACTCTAAATAAGGGAGAGCGTCG |

**Table S3:** List of the detected RNA targets (5’→3’) used in this study.

| HIV | GCUUGUACCAAUUGCUAUUGUAAAAAGUGUUGCUUUCAUUGCCAAGUUUGUUUCAUAACA |
| --- | --- |
| HCV | GCCUUGUGGUACUGCCUGAUAGGGUGCUUGCGAGUGCCCCGGGAGGUCUCGUAGACCGUGCAUCAUGAGCACAAAUCCUAAACCUC |
| P53 | CCCCUCCUGGCCCCUGUCAUCUUCUGUCCCUUCCCAGAAAACCUACCAGGGCAGCUACGGUUUCCGUCUGGGCUUCUUGCAUUCUGGGACAGCCAAGUCUGUGACUUGCACGUACUCCCCUGCCCUCAACAAGAUGUUUUGCCAACUGGC |
| SARS-CoV-2-N gene block | UCUGAUAAUGGACCCCAAAAUCAGCGAAAUGCACCCCGCAUUACGUUUGGUGGACCCUCAGAUUCAACUGGCAGUAACCAGAAUGGAGAACGCAGUGGGGCGCGAUCAAAACAACGUCGGCCCCAAGGUUUACCCAAUAAUACUGCGUCUUGGUUCACCGCUCUCACUCAACAUGGCAAGGAAGACCUUAAAUUCCCUCGAGGACAAGGCGUUCCAAUUAACACCAAUAGCAGUCCAGAUGACCAAAUUGGCUACUACCGAAGAGCUACCAGACGAAUUCGUGGUGGUGACGGUAAAAUGAAAGAUCUCAGUCCAAGAUGGUAUUUCUACUACCUAGGAACUGGGCCAGAAGCUGGACUUCCCUAUGGUGCUAACAAAGACGGCAUCAUAUGGGUUGCAACUGAGGGAGCCUUGAAUACACCAAAAGAUCACAUUGGCACCCGCAAUCCUGCUAACAAUGCUGCAAUCGUGCUACAACUUCCUCAAGGAACAACAUUGCCAAAAGGCUUCUACGCAGAAGGGAGCAGAGGCGGCAGUCAAGCCUCUUCUCGUUCCUCAUCACGUAGUCGCAACAGUUCAAGAAAUUCAACUCCAGGCAGCAGUAGGGGAACUUCUCCUGCUAGAAUGGCUGGCAAUGGCGGUGAUGCUGCUCUUGCUUUGCUGCUGCUUGACAGAUUGAACCAGCUUGAGAGCAAAAUGUCUGGUAAAGGCCAACAACAACAAGGCCAAACUGUCACUAAGAAAUCUGCUGCUGAGGCUUCUAAGAAGCCUCGGCAAAAACGUACUGCCACUAAAGCAUACAAUGUAACACAAGCUUUCGGCAGACGUGGUCCAGAACAAACCCAAGGAAAUUUUGGGGACCAGGAACUAAUCAGACAAGGAACUGAUUACAAACAUUGGCCGCAAAUUGCACAAUUUGCCCCCAGCGCUUCAGCGUUCUUCGGAAUGUCGCGCAUUGGCAUGGAAG |
| miRNA21 | UAGCUUAUCAGACUGAUGUUGA |

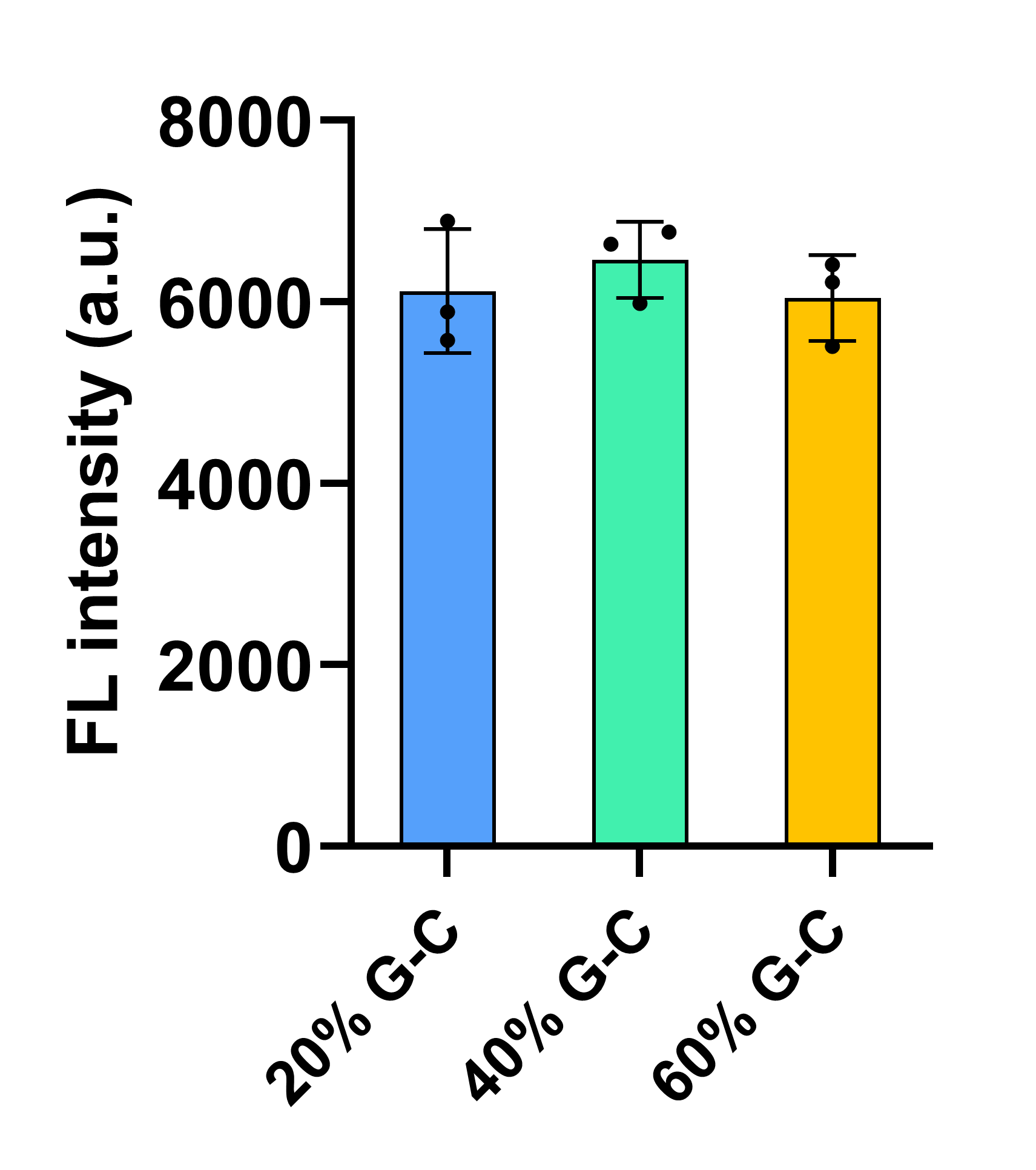

**Fig. S1. Investigation of the sequence dependence for S6 scaffold RNAs with varying G-C%.** The data demonstrate no apparent sequence-dependent effect; therefore, a S6 scaffold with 40% G-C content was selected.

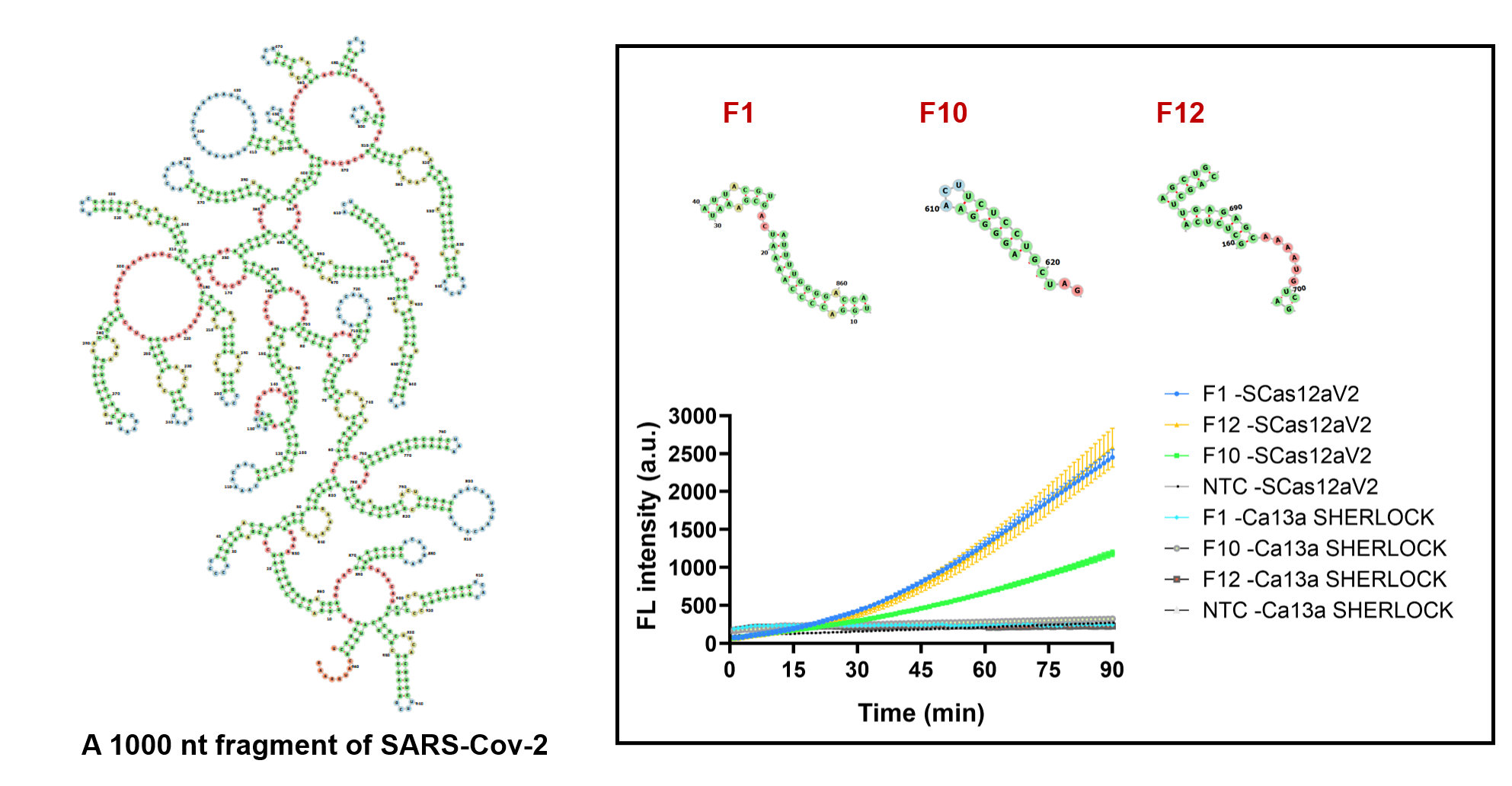

**Fig. S2. Detection of the same regions in a 1000-nt SARS-Cov-2 fragment by SCas12aV2 assay and SHERLOCK.** The findings demonstrate that the SCas12aV2 assay effectively detects highly structured RNA targets, which are not detectable by the Cas13a-based SHERLOCK method.

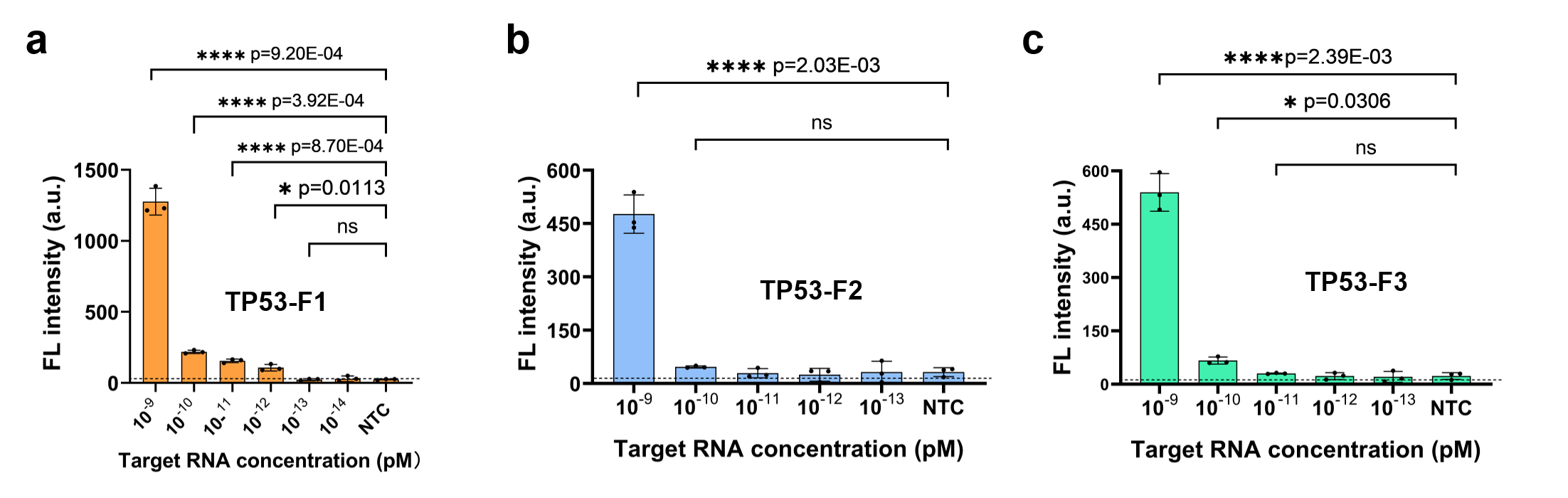

**Fig. S3. Determination of the limit of detection of the SCas12aV2 assay for RNA detection.** **a-c** Limit of detection of three TP53 targets was determined by a fluorescence assay. The plot illustrates the background-subtracted fluorescence intensity at t = 60 minutes for varying concentrations of the target. All the experiments were conducted in triplicate and error bars represent mean value +/− SD (n =3), and statistical analysis was conducted using a two-tailed t-test. Statistical significance was determined as follows: ns (not significant) for p > 0.05, * for p ≤ 0.05, ** for p ≤ 0.01, *** for p ≤ 0.001, and **** for p ≤ 0.0001.

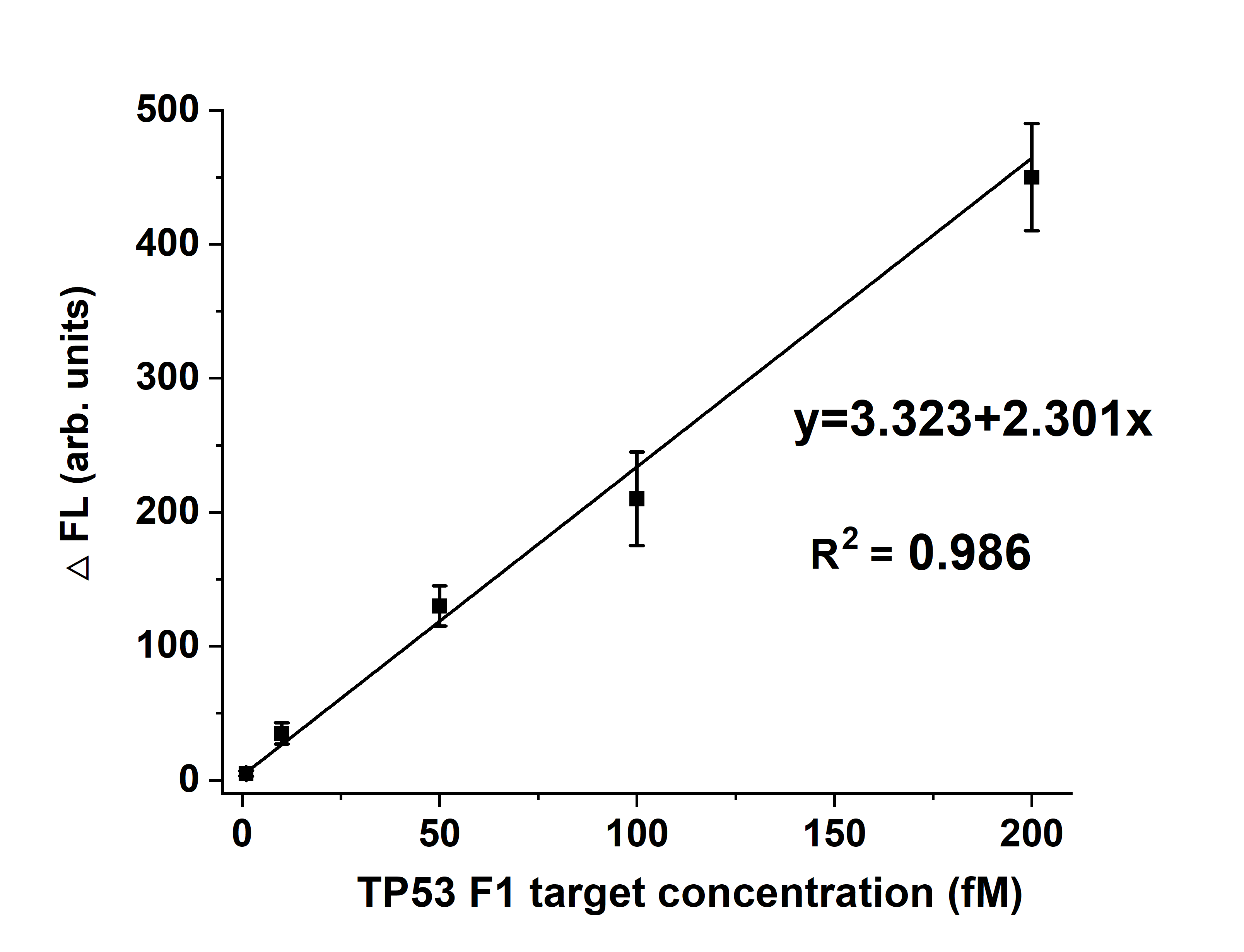

**Fig. S4.** **The linear relationship between fluorescence and TP53 F1 target concentration.** Error bars represent the standard derivation of three repetitive experiments. Source data are provided as a Source Data ﬁle.

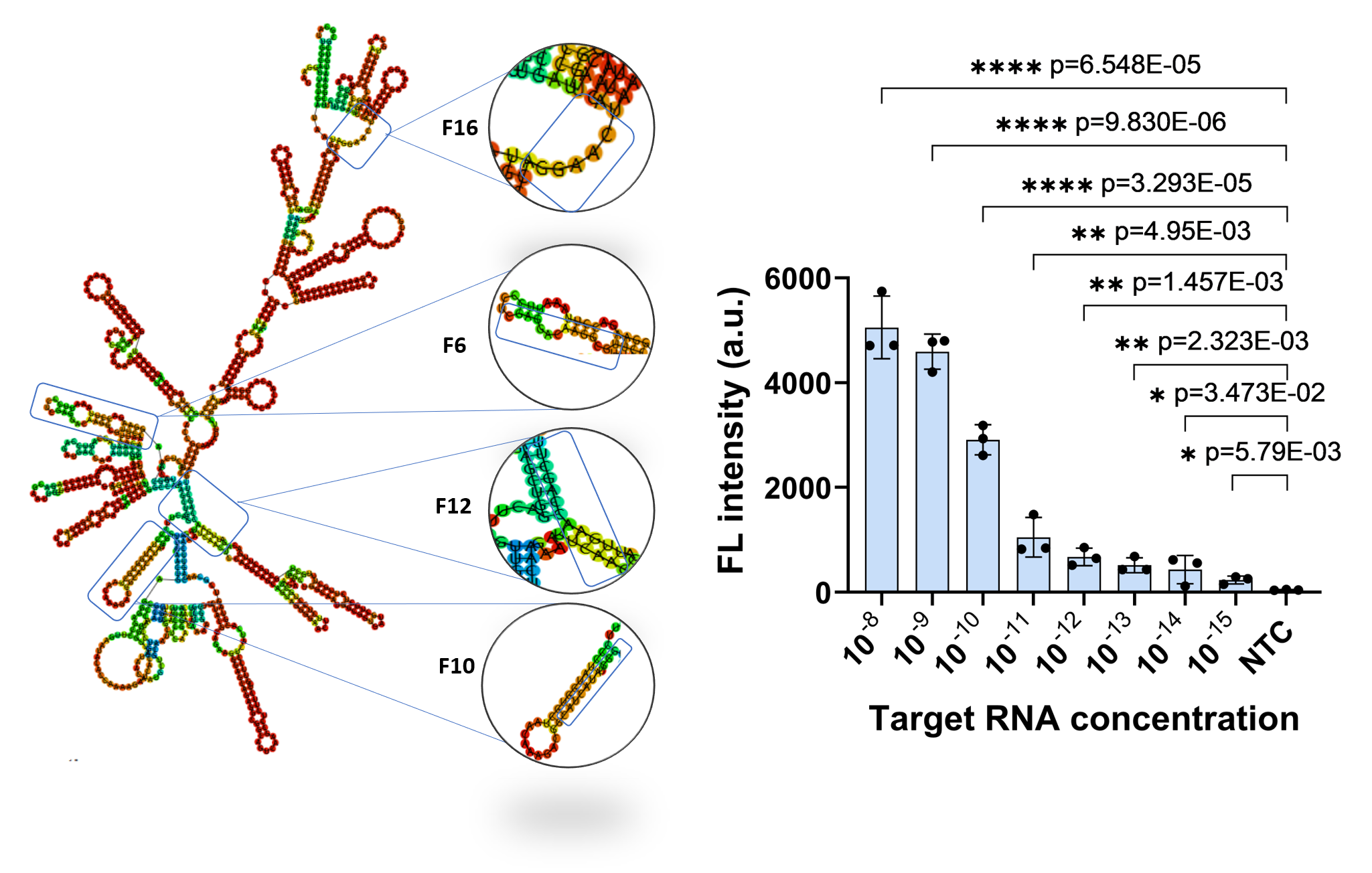

**Fig. S5.** **Determination of the limit of detection of the SCas12a fluorescence assay using pooled S6.1 activators.** The plot illustrates the background-subtracted fluorescence intensity at t = 60 minutes for varying concentrations of the target. Four distinct regions of a SARS-Cov-2 fragment were chosen as the targets and their corresponding S6.1 activators were employed. Error bars indicate the mean value ± standard deviation (n=3), and statistical analysis was conducted using a two-tailed t-test. Statistical significance was determined as follows: ns (not significant) for p > 0.05, * for p ≤ 0.05, ** for p ≤ 0.01, *** for p ≤ 0.001, and **** for p ≤ 0.0001. Source data are provided as a Source Data ﬁle.
